# Superior Intracellular Detection of Cytokines, Transcription Factors, and Phosphoproteins by CyTOF Compared with Fluorescent Cytometry

**DOI:** 10.1101/2025.05.30.656809

**Authors:** Michael J. Cohen, Erika L. Smith-Mahoney, Madison Bailey, Ling Wang, Lauren J. Tracey, Laura Polanco, David King, Christina Loh, Amedeo J. Cappione, Anna C. Belkina, Jennifer E. Snyder-Cappione

## Abstract

Unraveling biological complexity, whether it be immune subset distribution in infectious disease(s), autoimmunity or tumor heterogeneity, requires technologies capable of single-cell proteomic analysis, such as flow cytometry. Surface immunophenotyping alone is often insufficient, as interrogating functional capacity is required to determine cellular mechanisms and effectively inform diagnostic biomarker discovery, therapeutics and vaccine development. However, large panels with intracellular markers are subject to numerous challenges, including spectral overlap and background cellular autofluorescence, reducing resolving power for rare subsets or populations defined by low-abundance expression. We posited that mass cytometry may overcome such limitations; to address this, three small (11–12-plex) clone-matched antibody panels were evaluated by spectral flow and mass cytometry. Panels were comprised of surface and intracellular targets (phospho-epitopes, transcription factors or cytokines) and designed to minimize fluorescence spectral overlap. CyTOF™ technology offered superior signal resolution across the range of intracellular targets. Improved signal-to-noise provided better resolution of phospho-events and transcription factor expression, in particular TOX and T-bet. Most strikingly, stimulation-specific IL-10+ and IL-13+ cells were only detected by CyTOF. Superior resolution of these cytokines enabled accurate population clustering, permitting more unique immune cell signatures to be found, including Tr1 and Tc2 populations, thus providing a more comprehensive picture of the immuno-diversity present. Our findings indicate that CyTOF technology could catalyze seminal discoveries in functional immune profiling, driving therapeutic design and diagnostics.

## Introduction

High-dimensional single-cell analysis is a requisite tool for unraveling the complexity of the immune system and its role in homeostasis, disease pathologies and therapeutic interventions. In cancer, depending on the tissue of origin, intrinsic features of cancer cells, tumor stage and patient characteristics, the cellular composition and functional state of the immune response can vary widely and lead to a range of potential outcomes^1^. Measuring human cell functional signatures in a comprehensive, cross-lineage manner would provide key insights into several facets of immunology, including functional plasticity of innate T cell populations, T and B memory cell diversity following infection and/or vaccination, and the panoply of off-target (immunosuppressive, inflammatory) activities associated with cell therapies. While single-cell transcriptomics methods offer massive plexing capacity with wide dynamic ranges for variable signal detection, the correlation between RNA and protein expression is limited and thereby results are often misleading, including in the context of elements that modulate function in a temporal fashion (Sue et al. 2024). However, phenotyping of surface receptor signatures is insufficient and inconclusive to determine cell function; intracellular profiling of functional markers, such as cytokines, is required to accurately assess a cell’s state and functional potential.

Flow cytometry, as a highly sensitive, multi-parameter technology, has been a cornerstone for evaluating phenotypic diversity at the single-cell level through surface marker delineation and subset definition among immune cells in healthy and diseased individuals. Full spectrum flow cytometers (FSFC) are a recent advancement in instrumentation that measure the entire fluorescent emission spectrum across multiple lasers, thus permitting a unique spectral signature to be defined for each fluorochrome. Algorithms unmix the spectrally overlapping profiles, enabling far greater plexing capacity^2,3^. Still, the fluorochromes that possess wide emission spectra or share high spectral similarity, when combined in a multiplex panel, can outspan the unmixing capacity of the algorithm and lead to spreading errors in resultant data^4,5^. Mass cytometry is an orthogonal method of high-dimensional single-cell analysis that uses antibodies conjugated to heavy metal isotopes. In contrast to FSFC, CyTOF detects time-of-flight (TOF) for atoms derived from each of the metal tags; each atom’s TOF is a unique signature determined by mass, thus permitting determination of the atomic composition of each label for quantitative analysis of expression at the single-cell level ^6^. Three major differences with FSFC are key to the application of CyTOF technology to high-dimensional studies: (1) metal tags are exogenous cellular elements, therefore no background exists in samples, (2) signals derived from the discrete metal isotopes possess little to no overlap, and (3) metal tags are extremely stable and unhindered by the *ex vivo* conditions that can affect fluorophores, such as signal degradation over time, light exposure, temperature sensitivity and reactive oxygen species. These factors may contribute to mismatches in reference spectra, potentially leading to inaccuracies in unmixing, signal spreading errors and potential masking of dim signals. The CyTOF platform has provided detailed insights to advance discovery in the fields of cancer^7–9^, allergy/autoimmunity^10,11^, immune cell lineage and subsetting^12,13^, and anti-viral T cell responses^14,15^.

To date, several studies have compared the performance of FSFC vs. CyTOF technology for immune cell monitoring with an overall good concordance (correlation coefficient >0.98)^16–19^. While panels ranged up to 40 targets, these efforts focused primarily on surface markers and samples that were not cultured *ex vivo* or re-stimulated. Many intracellular functional readouts, including human Th2/Treg cytokines (IL-5, IL-10 and IL-13) and phospho-activation states, have been historically difficult to reproducibly detect by FSFC even with small panels and single cytokine measurements. Based upon these findings, we posited that CyTOF technology may overcome the limitations of FSFC in those applications, enabling better signal resolution for intracellular markers.

To address this question, we performed, to our knowledge, the first direct comparison of the two platforms using panels targeting multiple classes of intracellular markers. Our approach was designed to challenge the performance capacity while minimizing the potential bias or impact of spectral overlap and commercial availability of reagents on panel design and performance. We developed 3 small panels (11–12-plex) each with surface and intracellular markers, the latter targeting unique classes of functional readouts (phospho-epitopes, transcription factors or cytokines). All panels were designed with clone-matched antibodies and employed fluorescent conjugates that minimized spectral overlap. Donor PBMC were cultured, divided equally and stained with either the fluorochrome-or metal-conjugated antibodies (surface then intracellular) and acquired on a Cytek Aurora spectral cytometer or the CyTOF XT mass cytometer, respectively. Data sets from the cytokine panel were projected with opt-SNE to visualize functional cellular diversity. Overall, data collected by CyTOF demonstrated superior signal resolution for a range of intracellular readouts, including four of five phospho-activation states, the majority of transcription factors analyzed, and cytokines IL-10 and IL-13. The superior signal resolution achieved with CyTOF technology translated to the identification of more diverse functional subsets.

## Methods

### Human PBMC

For all three staining protocols, human PBMC from 8 healthy donors were obtained from STEMCELL Technologies. Donors were randomly selected and ranged in age (24–48), varied in gender, ethnicity, smoker or non-smoker, and CMV sero-status. For each independent experiment, cryopreserved PBMC from 4 donors were thawed in complete RPMI 1640 with 10% fetal calf serum (cRPMI) medium (Thermo Fisher Scientific) containing Anti-Aggregate Wash Supplement (Cellular Technology Ltd.), following manufacturer instructions.

### Panel development

Antibody clones were selected from commercial vendors based on literature review and availability for both platforms. All antibodies were clone-matched between mass and fluorescent cytometry panels. Lineage marker antibodies were titrated in the presence of live-dead discrimination reagent. Intracellular target antibodies were titrated in the presence of previously titrated lineage antibody cocktail and live-dead discrimination reagent. Titers were selected to ensure best possible signal resolution: separation from negative background in stimulation-specific targets and separation from fluorescence or metal minus one background for stimulation-independent targets.

### Phospho-protein staining

Thawed PBMC were counted and resuspended in cRPMI at a concentration of 2×10^6^ cells/mL and rested overnight at 37°C. Following resting, cells were harvested, counted, washed with PBS, and split evenly for FSFC and CyTOF staining. Cells were plated in 96-well v-bottom plates at 3×10^6^ cells/well. For single-stain reference controls required for FSFC, cells from all 4 donors were combined and plated onto corresponding wells.

For FSFC, cells were stained with LIVE/DEAD blue viability dye (Thermo Fisher Scientific), washed with FACS buffer and incubated with Fc-receptor- and monocyte-blocking solutions (BioLegend). Cells were then incubated with the surface antibody cocktail. For CyTOF, cells were washed with Maxpar™ Cell Staining Buffer (CSB) (Standard BioTools) and incubated with Fc-receptor-blocking solution (BioLegend). Cells were then incubated with the surface antibody cocktail in the presence of Cell-ID™ 103Rh Intercalator (Standard BioTools) for viability staining. Cells from both staining workflows were washed, resuspended in cRPMI and rested at 37 °C for 20 minutes. Cells were then stimulated with PMA (MilliporeSigma), IFNγ (BioLegend) and IFNα (BioLegend) at a final concentration of 100 ng/mL for each reagent. Unstimulated cells were left untreated. Samples were incubated at 37 °C for 15 minutes followed by immediate fixation to stop the stimulation. For FSFC, cells were permeabilized with BD Phosflow Perm Buffer III (BD Biosciences) while CyTOF samples were permeabilized with methanol. Samples were incubated on ice, washed and stained with the corresponding intracellular antibody panels. Following intracellular staining, samples were washed with FACS buffer or CSB for FSFC and CyTOF, respectively. CyTOF samples were fixed with 1.6% formaldehyde solution in PBS followed by resuspension in Maxpar Fix and Perm Buffer supplemented with Cell ID-Intercalator Ir (Standard BioTools). Samples from both workflows were stored at 2–8 °C overnight prior to acquisition.

### Transcription factor staining

Thawed PBMC were counted, washed in PBS, and divided evenly for FSFC and CyTOF staining. Cells were plated in 96-well v-bottom plates at 3×10^6^ cells/well. For single-stain reference controls required for FSFC, cells from all 4 donors were combined and plated onto corresponding wells. For FSFC, cells were stained with Zombie NIR viability dye (BioLegend), washed with FACS buffer, and incubated with Human TruStain FcX Fc-receptor blocking solution (BioLegend). Cells were then incubated with the surface antibody cocktail in the presence of monocyte blocker (BioLegend) and Brilliant Stain Buffer Plus (BD Biosciences). For CyTOF, cells were washed with CSB (Standard BioTools) and incubated with Fc-receptor blocking solution (BioLegend). Cells were then incubated with the surface antibody cocktail in the presence of Cell-ID 103Rh intercalator (Standard BioTools) for viability staining. After washing, samples from both cytometry workflows were fixed and permeabilized using the eBioscience Transcription Factor Buffer Set (Thermo Fisher Scientific). Samples were then incubated with the corresponding intracellular antibody panels followed by washes with FACS buffer or CSB for FSFC and CyTOF, respectively. CyTOF samples were fixed with 1.6% formaldehyde solution in PBS followed by resuspension in Maxpar Fix and Perm Buffer supplemented with Cell ID-Intercalator Ir (Standard BioTools). Samples from both workflows were stored at 2–8 °C overnight prior to acquisition.

### Intracellular cytokine staining

Thawed PBMC were counted and resuspended in cRPMI at a concentration of 2×10^6^ cells/mL and rested for 3 hours at 37 °C. Following resting, cells were harvested, counted and resuspended in cRPMI at a concentration of 5×10^6^ cells/mL. Cells were stimulated with phorbol 12-myristate 13-acetate (PMA) + ionomycin (BioLegend Cell Activation Cocktail, 1:1000 dilution) and lipopolysaccharide (LPS) (MilliporeSigma, 1 µg/mL). Unstimulated cells were left untreated. Immediately after the addition of the stimulation reagents, samples were seeded onto 6-well plates and incubated at 37 °C for 1 hour. Brefeldin A and monensin (BioLegend, 1:1000 dilution) were added to each well and samples were incubated at 37 °C for an additional 18 hours. Following the stimulation timeframe, cells were harvested, counted, washed with PBS, and evenly split for FSFC and CyTOF staining. Cells were plated on 96-well v-bottom plates at 3×10^6^ cells/well. For single-stain reference controls required for FSFC, cells from all 4 donors were combined and plated onto corresponding wells.

For FSFC, cells were stained with Zombie NIR viability dye (BioLegend), washed with FACS buffer and incubated with Human TruStain FcX Fc-receptor blocking solution (BioLegend). Cells were then incubated with the surface antibody cocktail in the presence of Monocyte Blocker (BioLegend) and Brilliant Stain Buffer Plus (BD Biosciences). After washing with FACS buffer, cells were fixed with eBioscience Intracellular Fixation Buffer (Thermo Fisher Scientific), permeabilized with eBioscience Permeabilization Buffer (Thermo Fisher Scientific) and incubated with the intracellular antibody cocktail. Cells were washed with FACS buffer and stored at 2–8 °C overnight prior to acquisition.

For CyTOF, cells were washed with CSB (Standard BioTools) and incubated with Fc-receptor blocking solution (BioLegend). Cells were then incubated with the surface antibody cocktail in the presence of Cell-ID 103Rh Intercalator (Standard BioTools) for viability staining. After washing with CSB, cells were fixed with Maxpar FIX-I Buffer (Standard BioTools), permeabilized with Maxpar Perm-S Buffer (Standard BioTools) and blocked with sodium heparin blocking solution (Sigma) in Maxpar Perm-S Buffer. Cells were then incubated with the intracellular antibody cocktail, washed with CSB and fixed with 1.6% formaldehyde solution in PBS. Cells were resuspended in Maxpar Fix and Perm buffer supplemented with Cell ID-Intercalator Ir (Standard Biotools) and stored at 2–8 °C overnight prior to acquisition.

### Sample acquisition and data analysis

FSFC samples were acquired on a Cytek Aurora 5-laser spectral analyzer and unmixed using SpectroFlo software version 3.2 (Cytek Biosciences) at the York University Flow Cytometry Facility. Single stain reference controls were prepared from the same cell cultures. Alternatively, UltraComp eBeads Plus Compensation Beads (Thermo Fisher Scientific) were used. CyTOF samples were acquired on a CyTOF XT system and samples were normalized with EQ™ Six Element Calibration Beads (Standard BioTools) using CyTOF software version 8.1. All FCS files were analyzed using FlowJo (BD Biosciences) to perform expert gating of the datasets for intracellular cytokine and transcription factor staining experiments. Cytobank cloud platform (Beckman Coulter) was used for phospho-protein staining experiments. OMIQ (Dotmatics) was used for automated computational analyses of the CyTOF cytokine dataset. Datapoints from equal numbers of live singlet lymphocyte events from 4 donors were concatenated and measurements from CD3, CD4, CD20, CD56, IFN-γ, IL-5, IL-4, IL-10 and IL-13 channels were projected into opt-SNE space^20^. Signal intensity of cytokine measurements was visualized as dotplot heatmap overlays. All experiments were repeated twice for each panel on two different days.

### ELISpot assays

Coating antibodies (anti-IL-4, -5, -10, -13, -IFN-γ, MabTech) were diluted to a final concentration of 10µg/mL using PBS. Biotinylated antibodies (anti-IL-4, -5, -10, -13, -IFN-γ, MabTech) were diluted in PBS 0.1%/Tween 20 + 2% BSA (PBS-TB) (R&D Systems). ELISpot plates (MilliporeSigma) were coated with primary antibodies for a minimum of 1 hour at room temperature. Plates were washed twice with PBS and once with RPMI 1640 Medium (Thermo Fisher) with 1% Gibco Penicillin-Streptomycin (Fisher) and 10% Corning Regular Fetal Bovine Serum (Thermo Fisher). Human PBMC (4 donors, STEMCELL Technologies) were seeded at 4 different concentrations (32.25K, 64.5K, 125K, 250K) in triplicate in complete media and cultured with (100 ng/mL PMA/2 mg/mL Ionomycin/2 mg/mL LPS) and without stimulation for 20 hours at 37 °C and 5% CO2. Plates were washed (2X PBS, 2X PBS+0.1% Tween 20, PBS-T). Plates were incubated with detection antibodies at room temperature for 30 minutes then washed 3 times with PBS-TB. Streptavidin-ALP (Mabtech) was added and incubated at room temperature for 45 minutes. Plates were washed twice with PBS-T, the plastic backings were removed and the plates were immersed in PBS-T for 1 hour. To develop spots, 100µL BCIP/NPT-plus (Thermo Fisher) was added and incubated at room temperature in the dark for 15 minutes. Each plate washed with DI water and 500 mL PBS then left to dry overnight. Spots were counted under a dissecting scope.

## Results

### Three low-parameter panels were designed to directly compare the intracellular detection capabilities of mass and fluorescent cytometry

Many factors can impact flow cytometric detection of signal, including spectral overlap, the antibody clone, and, in the case of functional detection, time and source of stimulation. In order to control for these confounding factors in our study, we: (1) designed small CyTOF and FSFC panels (the latter with minimal fluorescent spectral overlap); (2) used identical antibody clones in the CyTOF and FSFC panels to facilitate comparable reagent binding; and (3) utilized cells from cultures that were divided equally for CyTOF and FSFC, stained and run on the CyTOF or Aurora instruments, respectively. The 3 panels each included major lineage markers and detected phosphoproteins, transcription factors or cytokines from immune cells (**Table 1**). Importantly, for FSFC, the intracellular markers were assigned to the brightest channels possible, given reagent availability. Similarly, for CyTOF, mass channels within higher relative sensitivity were used for most intracellular markers. Finally, all antibodies were thoroughly titrated under the same cell culture conditions as the full panel experiments to ensure optimal signal to noise ratios for each marker.

**Table 1.** Three paired intracellular staining panels used to compare fluorescent and mass cytometry for the detection of cytokines, transcription factors and phosphoproteins.

| Cytokines |  |  |  | Transcription Factors |  |  |  | Phosphoproteins |  |  |  |
| --- | --- | --- | --- | --- | --- | --- | --- | --- | --- | --- | --- |
| Target | Clone | Fluorophore | Metal | Target | Clone | Fluorophore | Metal | Target | Clone | Fluorophore | Metal |
| CD3 | UCHT1 | BUV395 | 141Pr | CD3 | UCHT1 | BUV395 | 141Pr | CD3 | UCHT1 | BV570 | 141Pr |
| CD56 | NCAM16.2 | BUV737 | 176Yb | CD56 | NCAM16.2 | BUV737 | 176Yb | CD56 | R19-760 | BUV395 | 176Yb |
| CD8 | SK1 | BUV805 | 106Cd | CD8 | SK1 | BUV805 | 106Cd | CD8 | RPA-T8 | BUV805 | 162Dy |
| CD14 | M5E2 | APC-Fire 750 | 195Pt | CD14 | M5E2 | APC-Fire 750 | 195Pt | CD20 | H1 | BV510 | 144Nd |
| CD20 | H1 | BV510 | 144Nd | CD20 | H1 | BV510 | 144Nd | CD4 | SK3 | BV650 | 167Er |
| CD4 | SK3 | cFluor720 | 154Sm | CD4 | SK3 | R718 | 154Sm | pSTAT1 (Y701) | 4a | R718 | 153Eu |
| IL-5 | TRFK5 | APC | 167Er | Tbet | 4B10 | BV711 | 159Tb | pSTAT3 (Y705) | 4/pStat3 | AF647 | 158Gd |
| IFNγ | B27 | FITC | 116Cd | TOX | REA-473 | APC | 169Tm | pSTAT5 (Y694) | 47/Stat5 | BV421 | 147Sm |
| IL-4 | MP4-25D2 | PE-Cy7 | 172Yb | GATA3 | TWAJ | PE-Cy7 | 167Er | P38 (T180/Y182) | A16016A | PE | 165Ho |
| IL-10 | JES3-9D7 | BV421 | 165Ho | FOXP3 | PCH101 | PE | 162Dy | pERK1/2 (T202/Y204) | MILAN8R | AF488 | 171Yb |
| IL-13 | JES10-5A2 | BV711 | 169Tm | Live/Dead | N/A | Zombie NIR | 103Rh | Live/Dead | N/A | Blue | 103Rh |
| Live/Dead | N/A | Zombie NIR | 103Rh |  |  |  |  |  |  |  |  |

### CyTOF demonstrates superior resolution of stimulated phospho-activation states in CD4+ and CD8+ T cells

Historically, CyTOF technology has enabled effective multi-dimensional phospho-flow analysis across a range of experimental settings making CyTOF a platform of choice for intracellular signaling screening and fingerprinting^21–23^. However, no direct comparison of performance capabilities relative to conventional or spectral flow cytometry has been demonstrated to our knowledge. PBMC were surface stained, stimulated for 15 minutes in the presence of PMA, IFN-α and IFN-γ, then stained with phospho-specific antibodies (**Table 1**) and run on the respective instruments. Overall, the distinction between positive and negative events were clearly discernable for pSTAT1 from both platforms (**Figure 1A**). For pSTAT3, background is notably lower on the CyTOF plots, with bimodal gating not possible for CD4+ T cell data collected via FSFC due to data spread surpassing the positive signal shift (**Figure 1A**). Better separation of stimulated from unstimulated histograms was found for CyTOF vs FSFC for pSTAT5 and pERK1-2, while p38 signal was comparable between CyTOF and FSFC (**Figure 1A**). Quantification of these results revealed that CyTOF had higher phosphorylation signal over basal/unstimulated levels for all markers tested (**Figure 1B-C**).

**Figure 1.**
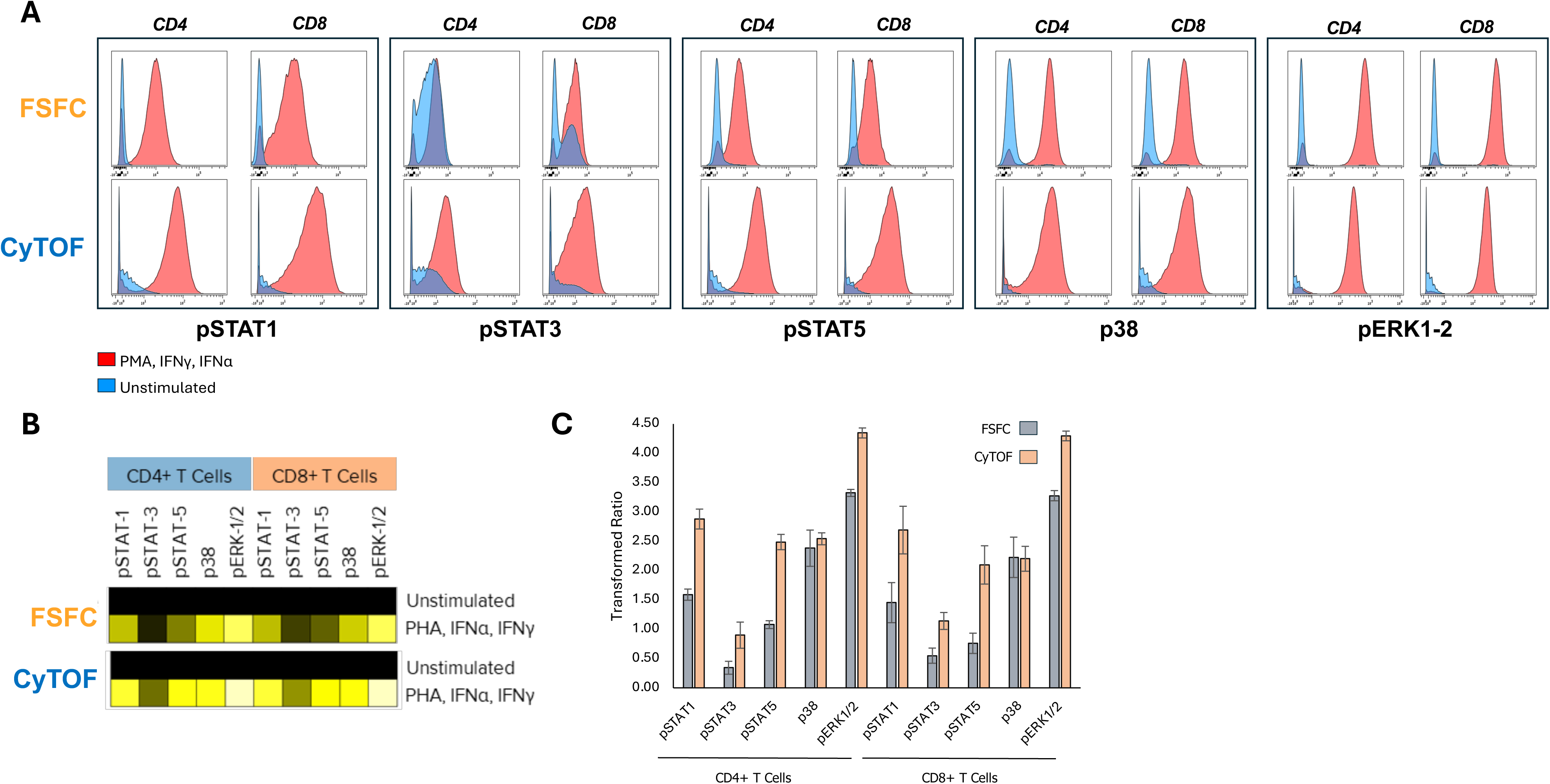
CyTOF demonstrates superior resolution of stimulated phospho-activation states. (A) Representative histogram overlays (unstimulated or PMA, IFN-α, IFN-γ-stimulated) from a single donor for 5 phospho-events (pERK1/2, p38, pSTAT1, pSTAT3, pSTAT5) in CD4+ and CD8+ T cells. (B) Heatmap depicts transformed ratios of median signal intensity values for protein phosphorylation compared to the unstimulated condition. (C) Bar graph depicts average transformed ratios of phosphorylation signal for each marker from 7 healthy donors (+/-S.E.M.).

### Superior detection of transcription factors (TF) using CyTOF as compared with FSFC

We next sought to determine if superior stimulation-specific intracellular detection of phosphoproteins was unique to these targets or if, alternatively, this improved signal extends to other types of intracellular readouts. To test this, we directly compared the detection of transcription factors using a clone-matched set of FSFC and CyTOF panels (**Table 1**). Cryopreserved PBMC samples were thawed and divided equally between the 2 cytometry workflows. In agreement with other reports, we found similar detection of FOXP3 with CyTOF and FSFC (**Figure 2A**). However, for other TFs, including T-bet and TOX, the signal resolution was superior in the CyTOF samples for both CD4+ and CD8+ T cells. Results from 2 TFs, TOX and T-bet revealed clear separation of double positive populations within T cells with CyTOF but not FSFC data (**Figure 2B**). Overall, the CyTOF data demonstrated superior signal:noise for transcription factors compared with fluorescent cytometry.

**Figure 2.**
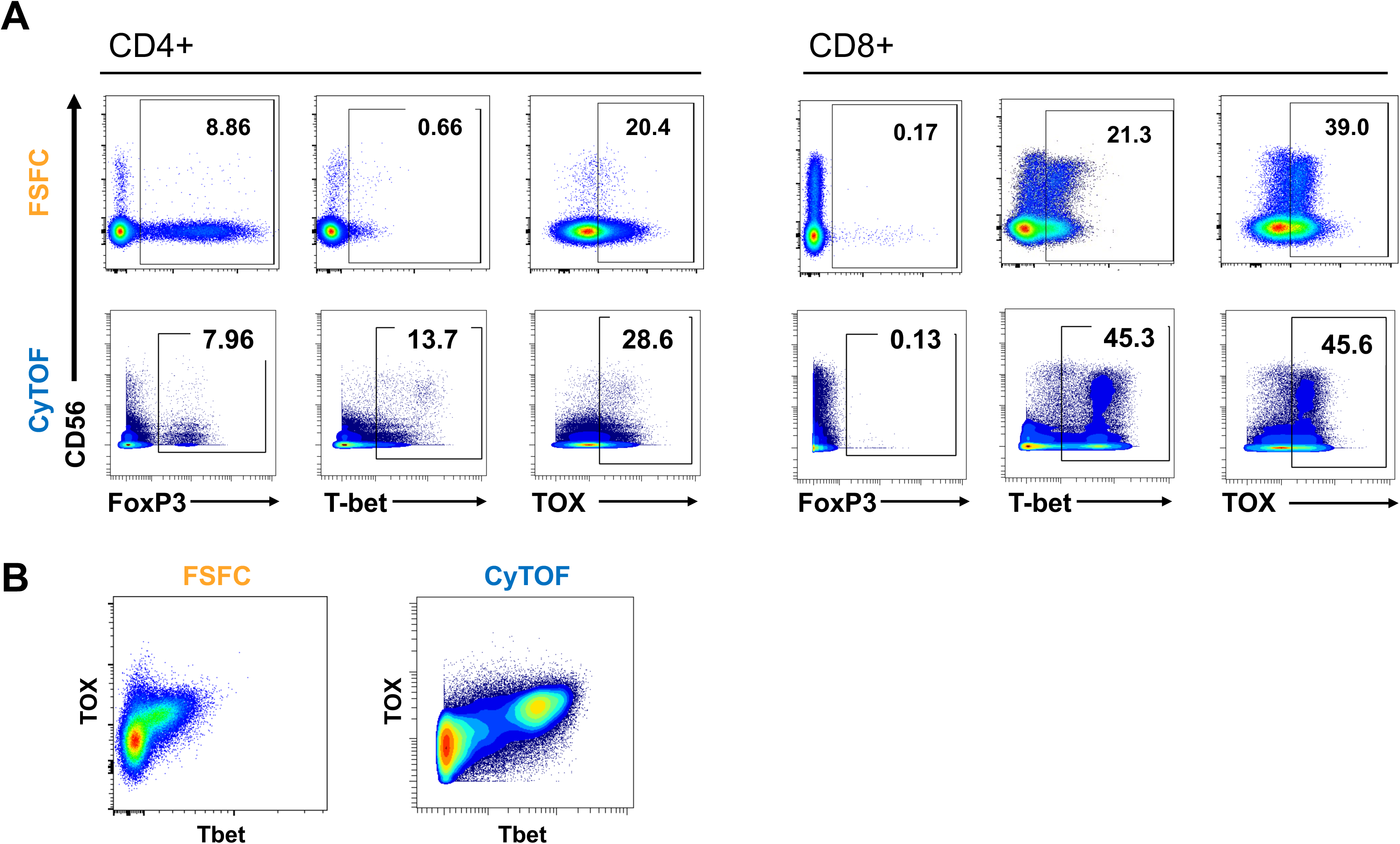
Overall higher signal-to-noise ratios for transcription factors using CyTOF as compared with FSFC. (A) CD4 and CD8 T cell subsets were defined; bi-variate plots show TF vs CD56 for each subset. Data represents the concatenation of 4 PBMC donors. Overall, FoxP3 reported very consistent frequencies between the two platforms for all subsets. By contrast, T-bet and TOX demonstrated higher CyTOF frequencies in both cases. (B) T-bet vs TOX bivariate plots (gated on CD3+ cells) demonstrating superior resolution of CyTOF.

### *Ex vivo*, stimulation-specific cytokine signal for IL-10 and IL-13 is clearly detected in human PBMC using mass but not fluorescent cytometry

We compared the cytokine detection capability of FSFC and CyTOF with the specific intention to include both cytokines that are commonly used in both platforms (IFN-γ, IL-4) and those historically more difficult to detect and/or less commonly reported in the literature from human samples (IL-10, IL-5, IL-13). PBMC were either stimulated with PMA/ionomycin and LPS or left untreated for 18 hours to generate samples with known positive cytokine signal and biological negative controls to establish assay background, respectively.

Direct comparison of IFN-γ and IL-4 expression from live T cells yielded overall similar results for both platforms in the appearance of events (distribution, separation of positive and negative signal (**Figure 3A**)) and the total percentage of positive events both with and without stimulation (**Figure 3C**). For IL-5, we observed similar results between the two platforms, with the exception of the presence of a small number of “bright” (high MFI) events in the FSFC unstimulated sample that was not seen with CyTOF (**Figure 3A**). However, IL-10 and IL-13 signal detection varied considerably between CyTOF and FSFC. Low frequency but clear, stimulation-specific IL-10+ T cell events were present in the CyTOF dataset **(Figure 3B)**. In contrast, the FSFC dataset showed a similar distribution of events with and without stimulation, with manual gating producing similar frequencies of positive events when guided by internal negative gating, making IL-10+ T cells non-detectable above assay background level (**Figure 3B**). This IL-10 finding was consistent across several donors (**Figure 3D**). In addition, and to our surprise, CyTOF detected a far greater frequency of IL-13+ CD4 T cells than FSFC post-stimulation across all donors (**Figure 3B**, **3D**). In the non-stimulated, negative control samples, IL-13 bright events were also found in the FSFC dataset but were not seen in the IL-13 non-stimulated result from CyTOF (**Figure 3B**) indicating higher background noise.

**Figure 3.**
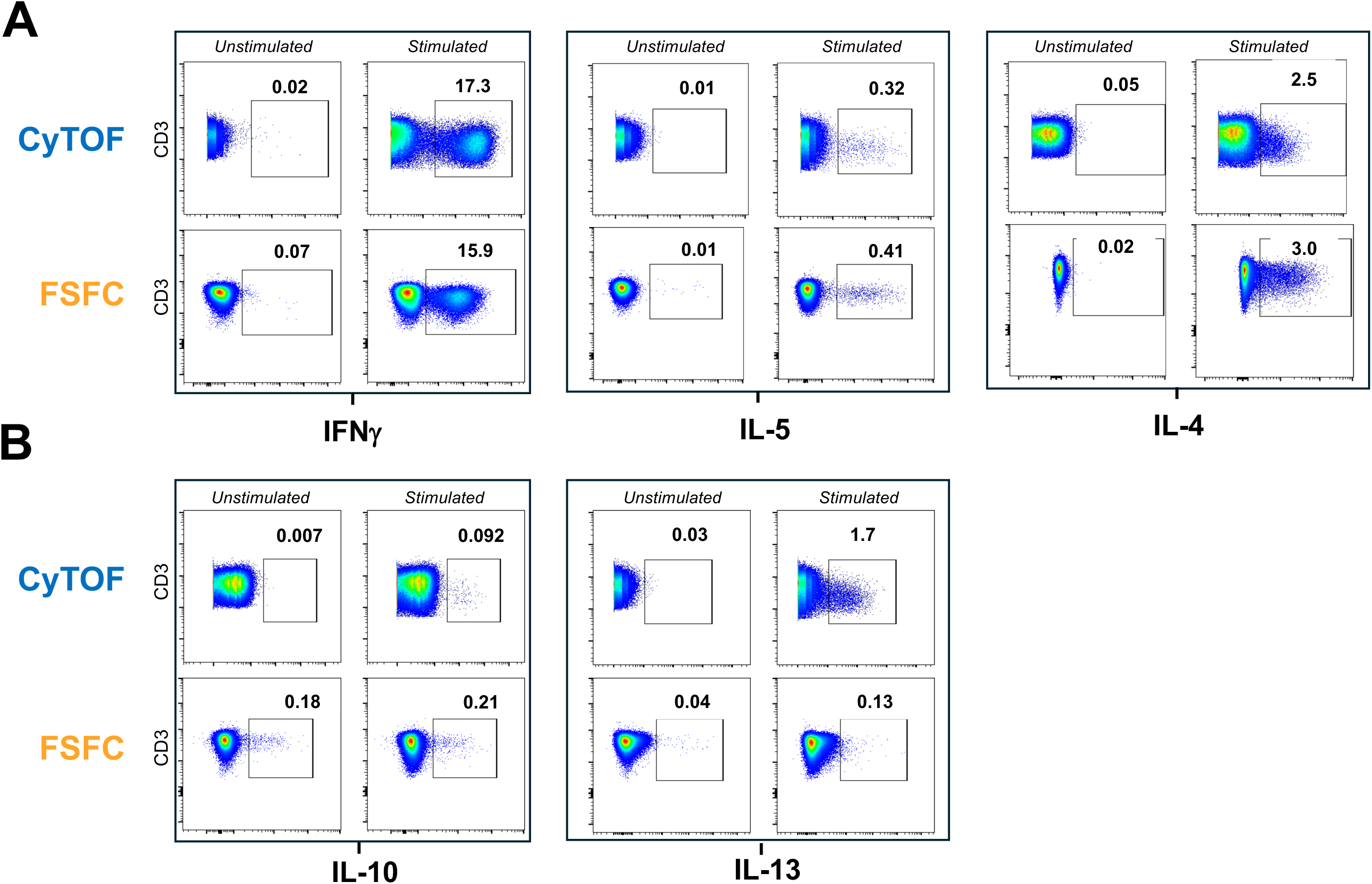
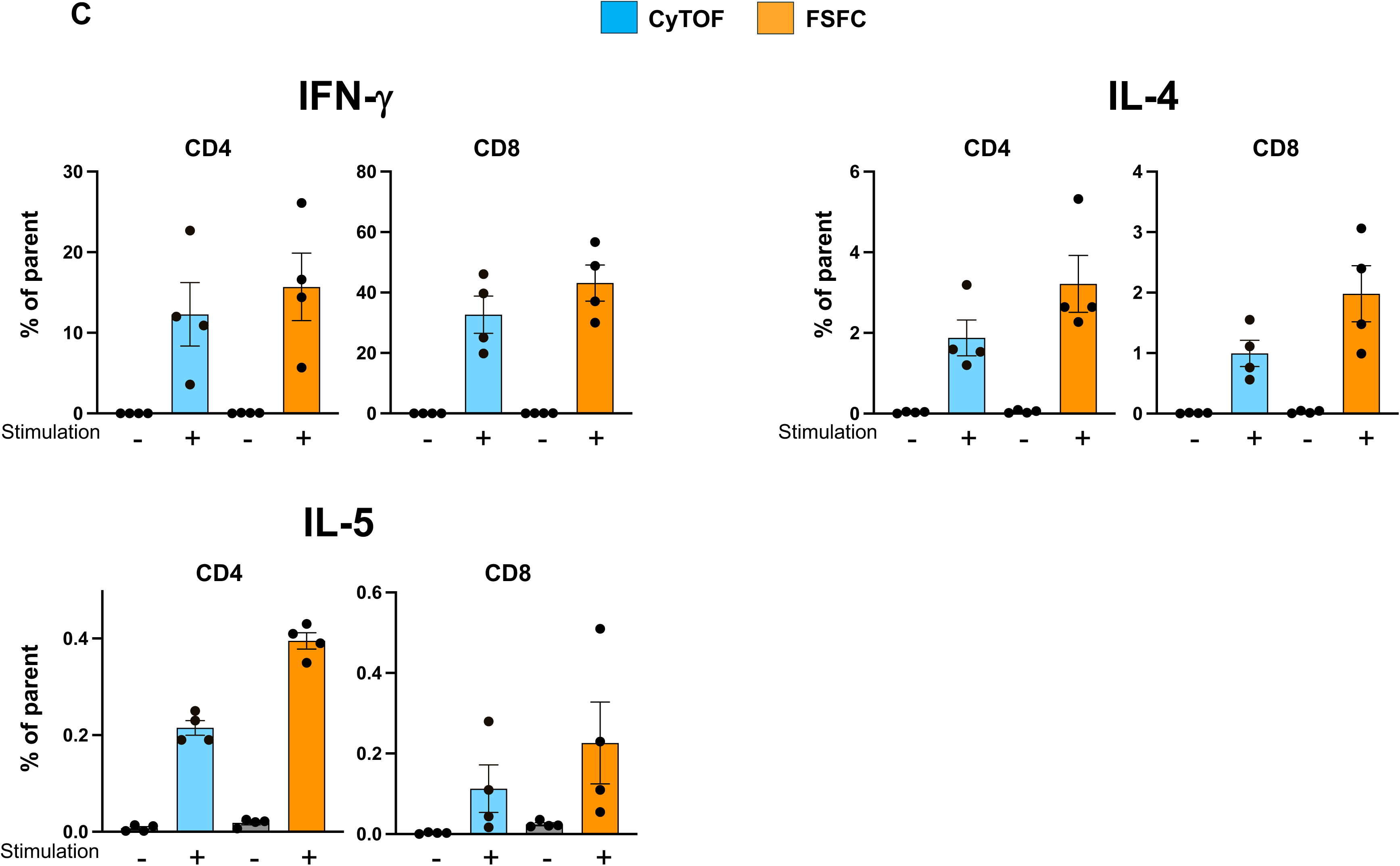
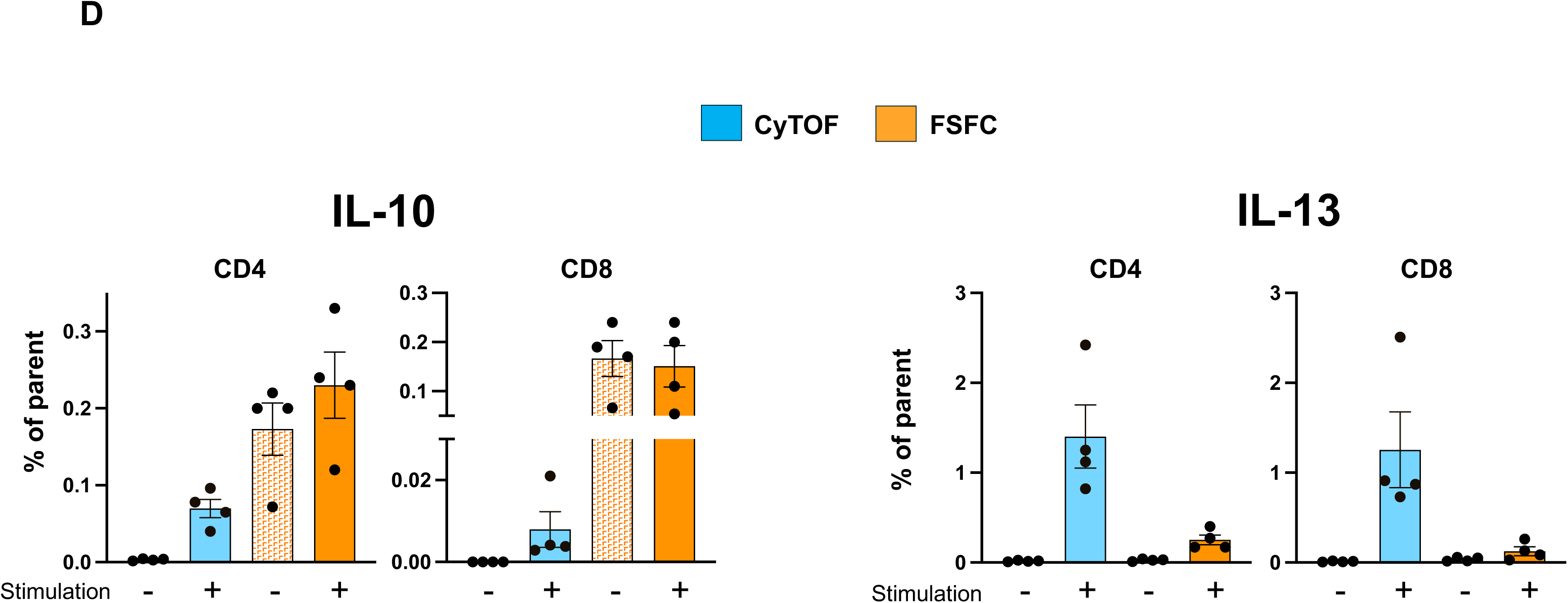
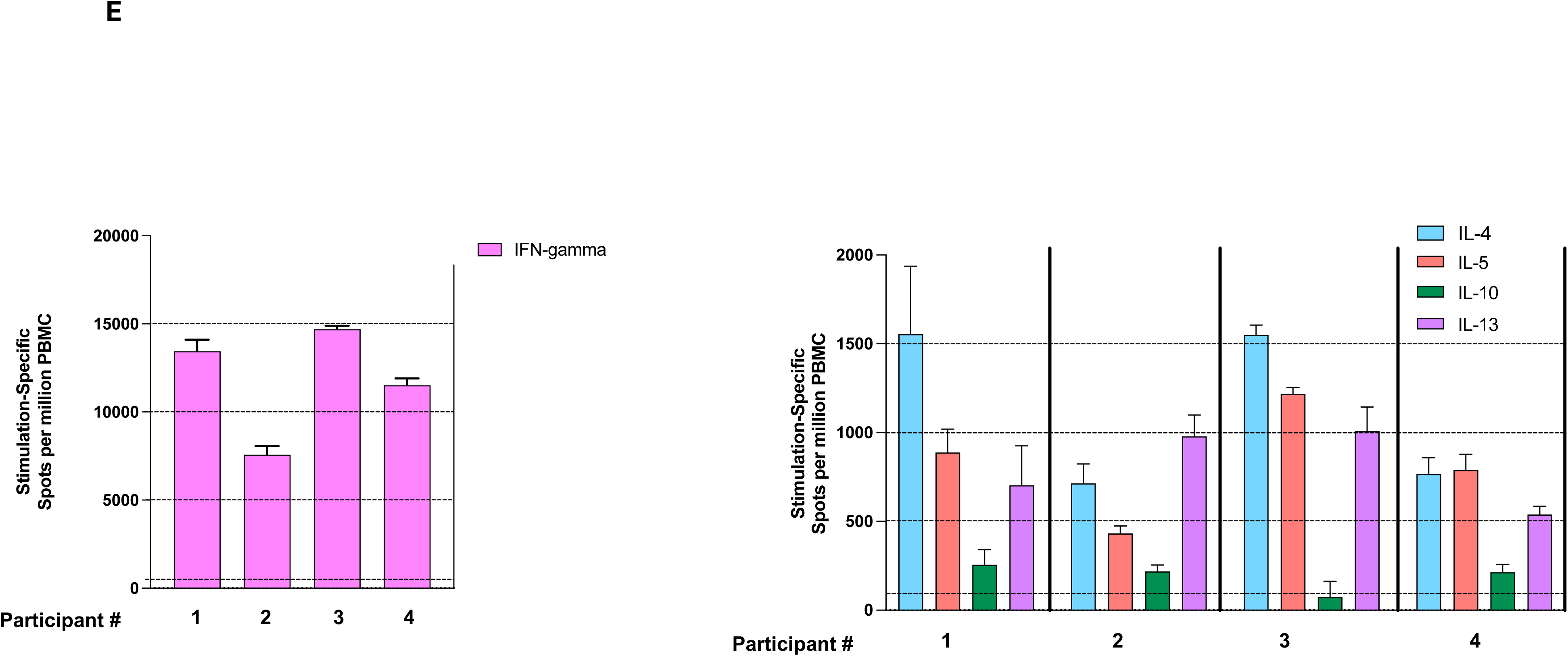
CyTOF enables clear detection of IL-10+ and IL-13+ human T cells in response to stimulation *ex vivo*. Plots show concatenated data from 4 healthy donors from unstimulated or PMA/ionomycin + LPS-stimulated conditions. An equal number of T cell events are displayed for the CyTOF and FSFC panels for each cytokine **(A, B)**. Percentages of the parent population (CD3+CD8-T cells) are shown above each gate. Data trends illustrated by the concatenated data shown above were consistent within each individual donor. IL-4 and IFNγ served as stimulation/assay positive controls. **(C, D)** Bar graphs depicting average % frequency of cytokine-producing CD4 or CD8 T cells from 4 donors (+/- S.E.M.); dots represent values for individual donors. **(E)** ELISpot assays of PBMC from 4 of the same donor PBMC lots included in A–D. Bar graphs depict spots per million PBMC of stimulated wells subtracted by signal of unstimulated wells. The values presented represent the mean spot number (per million cells) extracted from data points within the linear response curve for each effector function.

### Use of an orthogonal method (ELISpot) validates production of all 5 cytokines tested and agrees with CyTOF detection

To validate the detection of signal for IL-10 and IL-13 by CyTOF, we measured the frequencies of cytokine-producing cells from the same PBMC donors after a similar *ex vivo* stimulation using an orthogonal method. ELISpots for each of the 5 cytokines were performed on serial dilutions of total PBMC from the four donor samples (**Figure 3E**). Upon stimulation, all 5 effector functions were easily detected in each of the 4 donor PBMC at levels far above background. Stimulation-specific frequencies of cytokine producing cells in PBMC were similar between FSFC and CyTOF results for IFN-γ, IL-4 and IL-5. The frequencies of IL-10 and IL-13 secreting cells detected by ELISpot support the CyTOF but not the FSFC intracellular cytokine staining (ICS) results. Specifically, there was clear detection of stimulation-specific IL-10 secreting cells in a frequency consistent with the CyTOF dataset (**Figure 3D**). Similarly, IL-13 signal detected via ELISpot confirmed the presence of IL-13-secreting cells identified by CyTOF (**Figure 3D**). Though not directly comparable on a quantitative scale due to a number of differentiating factors (culture conditions, use of protein transport inhibitors, antibody clones, specific cell type analyzed), these ELISpot results confirm the presence of cytokine-secreting cells in PBMC at frequencies that parallel the CyTOF ICS results, thereby indicating that the signals detected by CyTOF are accurate and results from FSFC are markedly undercounting stimulation-specific IL10+ and IL-13+ cells.

### Co-expression of cytokines detected by CyTOF and visualized using opt-SNE validates accuracy of detected signals and reveals Tr1 and Tc2 populations

To investigate co-expression of the 5 cytokines measured with single cell resolution, we used opt-SNE^20^, a platform that uniquely enables clear visualization of rare populations within large datasets. We created an opt-SNE map of all collected PBMC events from 4 donors **(Figure 4, Center)**. Five major cellular islands were visualized, including CD4+ and CD8+ T cells, CD20+ B cells and CD56+ NK cells, as well as events negative for all T/NK/B cell markers in the panel **(Figure 4, Center)**. Heatmaps for each cytokine channel revealed their expression across the cellular landscape. Notably, small but clearly defined islands of IFN-γ+IL-10+ double producing and IL-10+ IFN-γ-CD4+ T cells were observed (**Figure 4A**, grouped as *A1* and *A2*, respectively). These populations were also clearly detected by bivariate plot visualization (**Figure 4B**). IFN-γ+ IL-10+ is a cytokine signature of Type 1 regulatory T cells (Tr1)^24,25^ (**Figure 4C**). Similarly, among CD8+ T cells, populations producing combinations of the type 2 cytokines IL-4, IL-5 and IL-13 were visualized (**Figure 4D**), including individual type 2 cytotoxic T cells (Tc2) that produce all 3 cytokines (**Figure 4E, F**). Tr1 and Tc2 cells were clearly enumerated in the samples; both populations are known to inhibit antigen presenting cell and/or T cell functions ^10,26^. It should be noted that opt-SNE maps created from FSFC data could not successfully resolve populations due to lack of signal in some channels (e.g., IL-13) and noise in the unstimulated conditions for others (e.g., IL-10) (data not shown).

**Figure 4.**
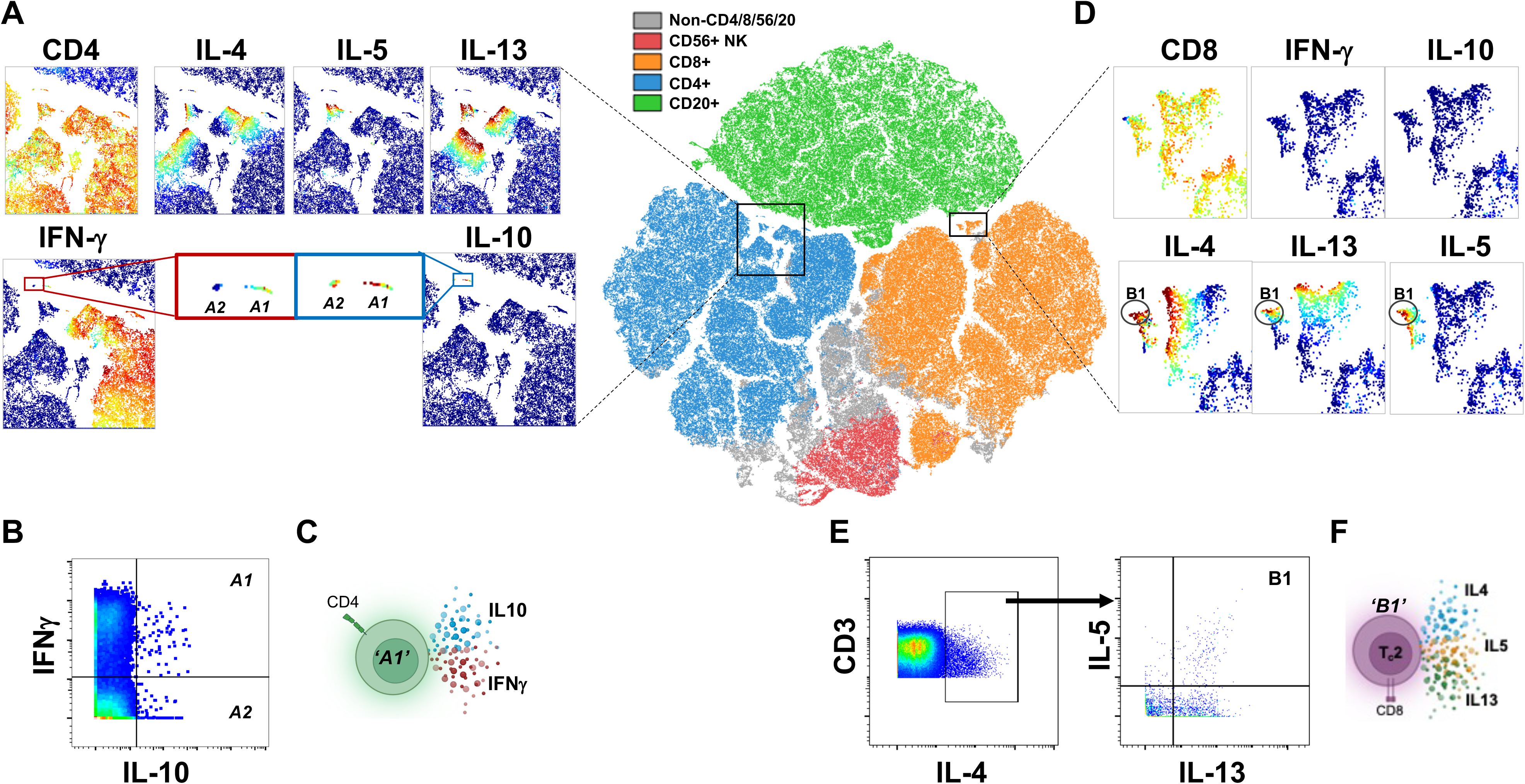
Tr1 and Tc2 populations are visible when applying opt-SNE to the CyTOF ICS dataset. (**Center**) Opt-SNE map of total PBMC concatenated from 4 healthy subjects. Based upon differential expression of CD4, CD8, CD20 and CD56, each color represents a unique surface marker-defined cell population. Data from equal numbers of live singlet lymphocyte events from 4 donors were projected into opt-SNE space. **(A, D)** Heatmaps and opt-SNE plot gradients (cold to warm) represent expression levels of the given antigen within each small cell cluster area (black squares) extracted from the whole opt-SNE map; warmer color signifies higher expression. Finer discrimination identified a small percent of T cells co-expressing IFNγ and IL-10 (“A1”, defined as Tr1 cells). **(B)** Bivariate plots confirm the presence of the double positive Tr1 cells. **(C)** Illustration of IL-10+IFNγ+ Tr1 cells. **(D)** Finer discrimination identified a small percent of triple-expressing IL4+IL5+IL13+ CD8+ cells (“B1”, defined as Tc2 cells). **(E)** Bivariate plots confirm the presence of IL-4+IL-5+IL-13+ Tc2 cells. **(F)** Illustration of IL-4+IL-5+IL-13+ Tc2 cells. Opt-SNE dimensionality reduction was informed by these cytokines permitting the unique multi-expressor phenotypes to be assessed within the cellular landscape mapped via other markers.

## Discussion

Cytometry by time in flight (CyTOF) and full spectrum flow cytometry (FSFC), as techniques capable of high-dimensional single cell analysis, have greatly enhanced the capacity to probe biological systems with high resolution ^27–30^. That said, each technology possesses strengths and weaknesses that act to inform and dictate their application ^17,19^. One area of expanding interest is determining functional potential of cells; this requires focus beyond surface marker detection and can optimally be ascertained through interrogation of multiple classes of intracellular readouts. To investigate relative performance, we executed, to our knowledge, the first direct head-to-head comparison of the intracellular detection capability of fluorescence and mass cytometry. Overall, the CyTOF-derived datasets demonstrated superior signal resolution compared to FSFC spanning all three of the intracellular readouts measured, including cytokines, transcription factors and phospho-activation events. Importantly, these results were found with low-parameter panels where every effort was made to minimize the impact of spectral overlap and autofluorescence on data processed from FSFC. These results are in stark contrast to the similar performance (detection, frequency, signal resolution) that has been seen between CyTOF and FSFC when comparative analyses were performed of surface markers in far larger panels^16–19^.

The improved signal-to-noise ratio of CyTOF data provided better resolution for phospho-activation events and transcription factor expression, in particular TOX and T-bet. Dysregulation of phospho-signaling pathways is core to the initiation and progression of malignancy. However, while being well established and validated for over 20 years^31–33^, phospho-flow has not demonstrated widespread use. Potential reasons for this include stringent culture conditions, variable performance in different fix/perm buffers, limited commercial availability of fluorescent conjugates, and poor or inconsistent signal detection by FSFC. CyTOF has previously demonstrated utility in studies of both transcription factors^19,34^ and phospho-signaling^8,9,35^.

Strikingly, within the cytokine panel, stimulation-specific IL-10- and IL-13-positive cells were only reliably detected by CyTOF. The validity of this finding was further substantiated through use of ELISpot assays; IL-10+ and IL-13+ PBMC were easily detected in all samples analyzed (Figure 3E). Cytokines of the Th2/Treg lineages, such as IL-5, IL-10 and IL-13, have been notoriously difficult to detect in human cells using fluorescence-based cytometry, to the point where the researchers are often compelled to combine IL-5, IL-13 and sometimes IL-10 readouts in the same fluorescence channel to boost the signal. A common practical solution to this issue is to switch the antibody-fluorochrome combination; however, we have tested multiple fluorochrome options and still were unable to recover IL-13 and IL-10 signal by FSFC (data not shown).The lack of reliable detection within these arms of the immune response has limited the understanding of their context and relative impact on the global immuno-signature (REF3). When the data sets were visualized with opt-SNE to determine functional diversity at the cellular level, the reliable detection of a greater number of anti-inflammatory functions with single-cell resolution by CyTOF translated to the specific identification of Tr1 and Tc2 subsets in human PBMC. It is the functional capacity of these rare T cell subsets that belies their significance as sources of immune defense or disease exacerbation, as well as their potential application as therapeutic agents. Given the functional signatures unmasked in this limited, small panel study, one can easily extrapolate the potential value of using CyTOF with far larger intracellular panels containing greater numbers of functional targets. With panel expansion, there is increasing incorporation of dimensionality reduction and clustering algorithms, such as opt-SNE, UMAP, FlowSOM and PhenoGraph; these tools reveal complex relationships within multi-dimensional data, thus enabling a better understanding of the cellular heterogeneity and identification of distinct cell populations^36–40^. However, algorithmic approaches require data of the highest quality to derive the most biologically relevant findings. For solution-based cytometry, this translates to a robust and reproducible assay (and platform) that permits distinct and quantifiable resolution of signal above background for each target. The data generated from the low-parameter cytokine panels in this study exemplifies this greatly. Due to the high resolution of low frequency cytokines such as IL-10 detected by CyTOF, the data set derived by CyTOF, when visualized with opt-SNE, reveals a wide breadth of functionally diverse cells within a population.

The cause of this difference in intracellular-resolving capacity between CyTOF and FSFC is unclear; however, several possibilities should be considered. FSFC analyzes a cell as a solid structure, whereas cells analyzed with CyTOF are atomized into an ion cloud and non-relevant atoms are eliminated prior to interrogation. In addition to differences in sensitivity between platforms, antibody size and hydrophobicity related to fluorochrome conjugations may impact performance. Moreover, our choices of small panels (11– 12 plex) and spectrally distinct fluorescent conjugates were made to minimize the potential impact of spillover spreading error (SSE) on the variance of unmixed signal^41–45^. Recent findings suggest that an additional level of variance, unmixing dependent spreading (UDS), imparted by the algorithmic correction of spillover, can negatively impact signal:noise, an effect that is independent of target expression and increases with panel size^46^.

CyTOF and FSFC possess unique strengths and weaknesses; thoughtful examination in platform choice is required to maximize data output while minimizing burden to perform the study. Beyond the performance differences outlined, CyTOF possesses strengths that offer clear benefits to a researcher exploring the use of larger panels (>25 markers) including functional readouts defined by intracellular staining. The issues with spectral overlap require use of single stains for efficient titration of each antibody and reference controls for each fluorophore during optimization and sample analysis, none of which are required for CyTOF. In addition, kit-based, in-house labeling of antibodies is commonplace with CyTOF, adding greater flexibility in panel development; FSFC suffers from variability in labeling performance, limited commercial availability and complexity in choice of chemistry that have prevented its routine application. Furthermore, the most optimal and brightest fluorophores were selected for the intracellular markers used in these low-parameter panels, given commercial availability of antibody conjugates. It can be appreciated that expanding the number of intracellular functional markers in the FSFC panels used in this study would be an increasingly difficult task, while CyTOF panel expansion would be more streamlined.

In summary, CyTOF technology provided a superior alternative for multiplex intracellular detection because it consistently demonstrated better positive signal above background. Future work will focus on development of a CyTOF-based 50-plus marker panel containing >20 intracellular functional markers of disease relevance. The panel will include cytokines spanning all known T cell functional lineages (Th1/Th2/Th9/Th17/Th22/Tfh/Treg) as well as targets linked to cell repair, chemoattraction and other cellular activities. Such insights, previously unseen, may better inform therapeutic decision-making, patient stratification and predictive outcomes^47–49^ through identification of biomarkers^9,50,51^. While the current study was limited to three small panels, the deeper, wider functional profiling enabled by CyTOF technology, when integrated with other omics data such as transcriptomics and metabolomics, provides a pathway to achieving the goals of personalized molecular insights and precision medicine.

